# Simultaneous assessment of eight phosphorylated STAT residues in T-cells by flow cytometry

**DOI:** 10.1101/2021.11.22.469156

**Authors:** Emily Monk, Melinda Vassallo, Paulo Burke, Jeffrey S. Weber, Pratip Chattopadhyay, David M. Woods

## Abstract

Signal transducer and activator of transcription (STAT) proteins are a family of transcription factors controlling functions in immune responses and other cell types. Given their importance, we developed a flow cytometry panel to assess eight phosphorylated STAT residues in human T-cells, including six tyrosine residues across six STAT proteins (STAT1, STAT2, STAT3, STAT4, STAT5a, STAT6) and additional serine residues on STAT1 and STAT3. We applied this protocol to test the *in vitro* induction of pSTATs in response to CD3/CD28 activation and a panel of recombinant cytokines. We also assessed the pSTAT expression profiles of naïve CD4+ T-cells polarized to Th1, Th2, Th17 or iTregs. pSTAT1(S727), pSTAT2(Y689) and pSTAT3(S727) were constitutively expressed in most T-cells, even in the absence of stimulation. For pSTAT1(S727) and pSTAT3(S727), we observed two positive states, high and low. Conversely, expression of pSTAT1(Y701), pSTAT3(Y705), pSTAT4(Y693) and pSTAT6(Y641) were absent in resting T-cells and only expressed with CD3/CD28 activation or with specific cytokines. Variable frequencies of pSTAT5a(Y694) expression were observed in resting T-cells, which increased with activation or specific cytokine stimulation (e.g. IL-2). IFNβ stimulation enhanced frequencies of expressing cells for all pSTATs. Correlations among several pSTATs, particularly pSTAT1(S727)^high^ and pSTAT3(S727)^high^ were observed. While polarization resulted in increases in canonically associated pSTATs, other non-canonical pSTAT changes were also observed. Collectively, we developed, optimized, and tested a sensitive and rapid approach for simultaneously assessing phosphorylation of six STAT proteins. Using this approach, we made several novel observations of T-cell pSTAT induction in response to stimuli.

## Introduction

Signal transducer and activator of transcription (STAT) proteins are a pleiotropic family of transcription factors controlling cellular functions in various cell types. There are seven STAT proteins in humans: STAT1, STAT2, STAT3, STAT4, STAT5a, STAT5b, and STAT6^1^. STAT5a and STAT5b are tandem genes with 91% residue similarity at the protein level; while some distinctions in function have been identified, most functions overlap^2–4^. STAT proteins enable cellular responses to external stimuli by communicating between transmembrane receptors and the nucleus. Cytokines binding to their respective receptors induces phosphorylation by Janus kinases (JAKs), which in turn phosphorylate STAT proteins. pSTATs then form hetero- or homodimers, translocate into the nucleus, and bind to associated DNA motifs initiating gene transcription programs.

STAT proteins are broadly involved in immune responses, including regulating immune cell proliferation, polarization, and memory formation^5–7^. In T-cells STAT1 signaling regulates the polarization and function of Th1 T-cells, STAT6 Th2 T-cells, STAT3 Th17 T-cells and STAT5 Tregs^8–11^. These phenotypes are induced by cytokine signaling including IL-12 (STAT1), IL-4 (STAT6), IL-6 (STAT3), and IL-2 (STAT5) ^3,8,11^. STAT3 also regulates the proliferation and memory formation of T-cells ^6^.

Several diseases have been associated with dysregulation of STAT proteins and the JAK-STAT signaling pathway. Many cancer studies have implicated various STAT proteins in tumor progression, antitumor immune responses, and patient response to immunotherapy^12–16^. For example, persistent STAT3 signaling in tumor cells is characteristic of most human cancers and is negatively associated with patient response to immunotherapy and overall survival^17–21^. Conversely, upregulation of T-cell pSTAT3 signaling was shown by our group to be associated with checkpoint immunotherapy response in metastatic melanoma^13^. Additionally, several autoimmune diseases have been associated with mutations or deregulation of the JAK-STAT pathway^22–26^.

Interrogating of all known STAT proteins simultaneously at single cell resolution is rarely performed. Further, it is rare for more than one residue to be assessed for a single STAT protein, despite phosphorylation of different residues conferring unique signaling and translocation characteristics^27–29^. In the present study, we have developed a flow cytometry panel to assess tyrosine phosphorylation of six STAT proteins (STAT1, STAT2, STAT3, STAT4, STAT5a, STAT6) and additional serine residue phosphorylations on STAT1 and STAT3.

## Materials & Methods

### T-cell Samples

Healthy donor PBMC were obtained from New York Blood Center (New York, NY USA) under an exempted IRB protocol at NYU Langone Health. T-cells were isolated with an EasySep CD3+ enrichment kit from StemCell Technologies (Vancouver, BC Canada), per the manufacturer’s protocol. For all experiments, isolated T-cells were cultured in X-VIVO media with 8% human AB serum on 96-well plates.

### Activation

For antibody activations, plates were coated with CD3 antibody (clone OKT3 at 0.5μg/mL) purchased from BioLegend (San Diego, CA USA) 24 hours prior to start of experiment. Unbound antibody was rinsed with PBS prior to plating of T-cells. Immediately after plating of T-cells, soluble CD28 antibody added (clone CD28.2 at 0.5μg/mL) (BioLegend). Other activation types assessed include: Dynabeads (1:1; 200,000/well) (ThermoFisher; Waltham, MA USA), Dynabeads (1:10; 20,000/well), or Immunocult Human CD3/CD28 T-cell Activator (5uL/well).

### Cytokine Stimulation

Kinetics were evaluated by adding recombinant cytokines to cultures at 24 hours, 2 hours, 1 hour, 30 minutes, or 15 minutes prior to fixing. The last 10 minutes of each time point included the addition of viability staining. Treatment wells were randomly spread throughout the plate to avoid plate location effects. All recombinant cytokines were obtained from R&D Systems (Minneapolis, MN USA).

### Polarization

Wells in a 96 plate were coated with CD3 antibody (OKT3 clone, 0.5ug/mL) 24 hours prior to start of experiment, and unbound antibody rinsed with PBS prior to plating of T-cells. Healthy donor naïve CD4+ T-cells were isolated with a StemCell EasySep naïve CD4+ isolation kit per the manufacturer’s protocol. Cells were then plated in the CD3 coated 96-well plate. Additional samples were cultured an uncoated plate as controls. Soluble CD28 antibody (CD28.2 clone, 0.5ug/mL) was added to all wells on the CD3 coated plate. Wells in each plate were treated with the following polarization mixes: None, Th1 – 1μg/mL IL-4 neutralizing antibody (clone 11B11), 5IU/mL IL-2 and 10ng/mL IL-12; Th2 – 1 μg/mL IFNγ neutralizing antibody (clone R4-6A2), 5IU/mL IL-2 and 10ng/mL IL-4; Th17 – 1μg/mL IFNγ neutralizing antibody (clone R4-6A2), 1μg/mL IL-4 neutralizing antibody (clone 11B11), 1μg/mL IL-2 neutralizing antibody (clone JES6-1A12), 1ng/mL TGFβ, 20ng/mL IL-6 and 50ng/mL IL-23; and inducible Tregs (iTreg) – 1μg/mL IFNγ neutralizing antibody (clone R4-6A2), 1μg/mL IL-4 neutralizing antibody (clone 11B11), 2ng/mL TGFβ and 300IU/mL IL-2. At the 30-minute timepoint, all cells in the uncoated plate were harvested. Three wells from each polarization in the coated plate were harvested at the 30 minute, 24-hour and 96-hour timepoints.

### Flow Cytometry

T-cells were viability stained with Live Dead Near IR amine reactive dye (ThermoFisher), fixed/permeabilized and antibody stained per the protocol included in the supplemental materials. Antibodies, clones and manufacturers used are shown in **Table 1**. Fluorescence minus one (FMO) controls were used to discriminate background fluorescence from positive signal. An Attune NxT (ThermoFisher) flow cytometer with blue, violet, yellow and red lasers (14-colors total) was used for data acquisition.

**Table 1.**
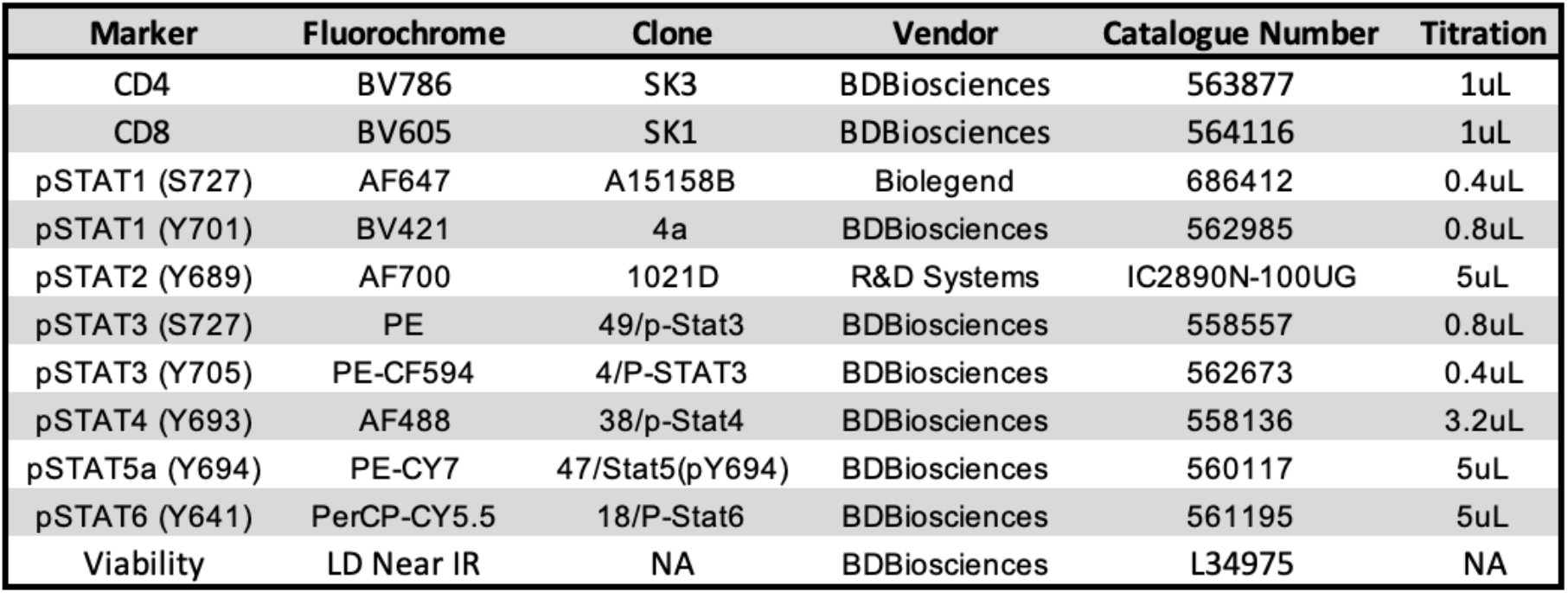
Flow cytometry panel design.

### Data Analysis

FlowJo v10.7 or v10.8 software (Ashland, OR USA) was used for initial analyses. Events were gated as shown in **Figure 2**. Phenograph^30^, UMAP^31^ and ClusterExplorer plugins for FlowJo were used for clustering and cluster composition analysis. Plugins were run on the concatenated fcs file; plugin values were extracted by gating samples in concatenated file by keyword gating. Data exported from FlowJo was analyzed with R v4.0^32^ using the R Studio interface. Flow fcs files were imported with the FlowCore package^33^. Data frames were manipulated using packages from the Tidyverse family^34^. Visualizations were produced using the ggplot2^35^, polychrome^36^, and ggpubr^37^ packages. For each experiment, both CD8+ and CD4+ T-cells were visualized. Given the high degree of similarity, in most figures data from CD4+ T-cells are shown as representative. In recombinant cytokine treatments experiments, significance was assessed by comparing each cytokine stimulation group against the untreated group by t-tests with a Bonferroni adjustment for multiple comparisons. For network graphs, pSTATs in CD4+ T-cells were assessed for correlations using the Pearson correlation coefficient and visualized as a network using the Hmisc^38^ and igraph^39^ packages. Network graphs for this manuscript were generated using Cytoscape software^40^. To improve visual interpretability, only edges with an associated p-value less than or equal to 0.05 were plotted.

### Rainbow & Cloud Plot

We generated custom visualizations of Phenograph cluster composition, which we term Rainbow & Cloud plots (**Figure 5D**). Graphs of the values of each channel for each Phenograph clusters were generated in R with the ggplot2 package. FMO values were down sampled to ease computational burden and plotted on top of channel values to discriminate values above background signal. The code for these visualizations and all others is included in the supplemental materials.

### Data Availability

Data frame csv files, fcs files, and R analysis code for all data analysis is included in the GitHub repository (https://github.com/SciOmicsLab/pSTAT_Data_Supplemental). R analysis code has been placed into a docker container to preserve package functionality. The GitHub repository includes instructions on how to use the docker container and run the analysis code.

## Results

### Cell viability staining can be accomplished rapidly

Manufacturer supplied protocols for amine reactive viability dyes require staining in PBS for 30 minutes, and DNA intercalating dyes (e.g. 4’6-diamidino-2-phenlyindole (DAPI)) are incompatible with fixation/permeabilization of cells needed for intracellular staining. However, phosphorylation of STAT proteins can occur rapidly and transiently in response to cytokine stimulation. Therefore, we first sought to determine the compatibility of viability staining in cell culture media and the minimal time necessary for viability staining of cells. Human T-cells were stained for 0-60 minutes, shown in **Figure 1**, by the addition of Live Dead Near IR amine reactive dye directly to the cell culture media, centrifuged, and resuspended in fixing buffer. Discrete populations of viable (Live Dead Near IR negative) and non-viable were discernable with as little as one minute of culturing with the dye. Importantly, these results indicated that viability staining could be done in cell culture conditions alongside short term stimulation (e.g. addition of cytokines).

**Figure 1.**
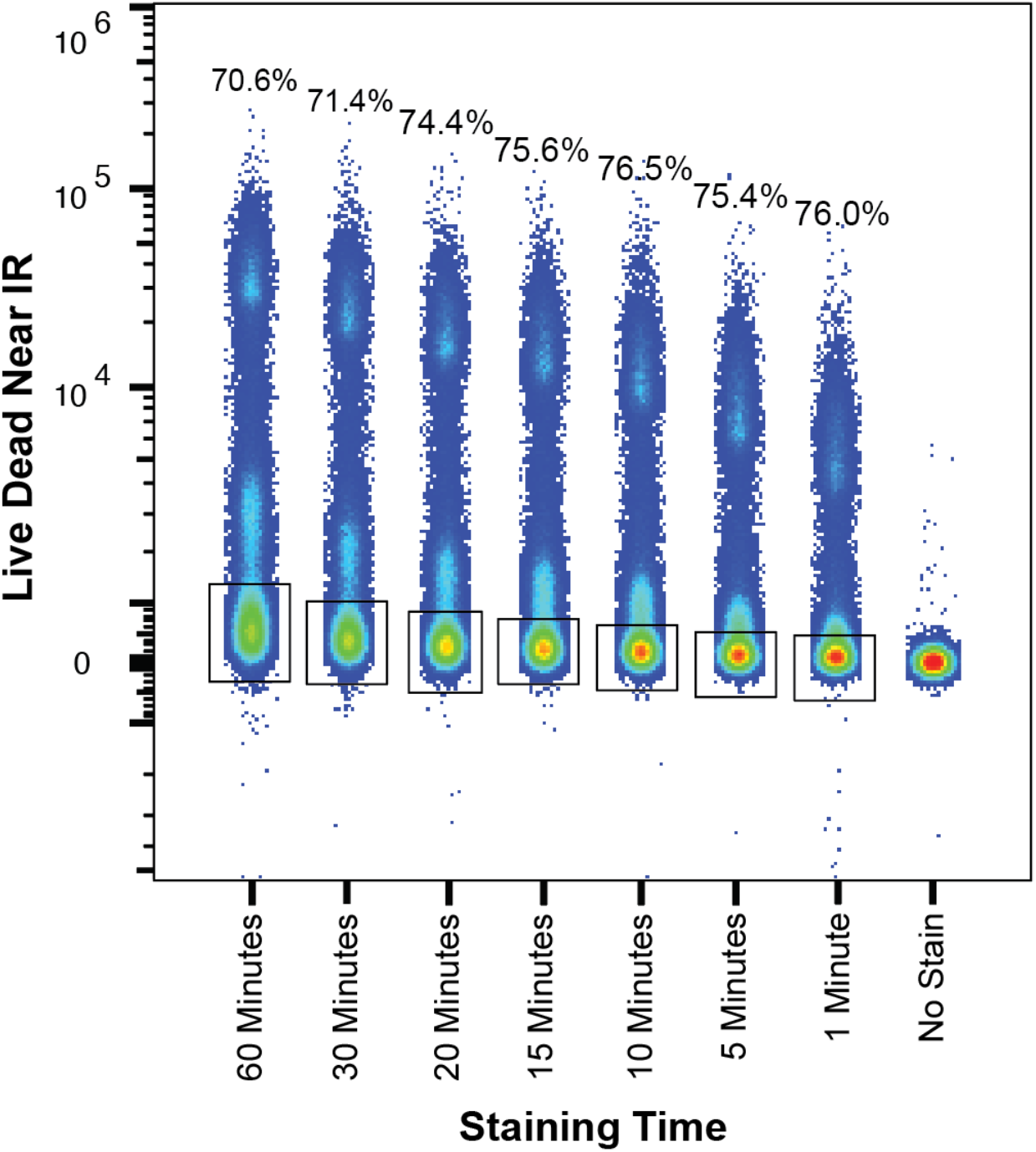
Minimal staining time with amine reactive viability dye is needed. PBMC were stained for the indicated times with the amine-reactive viability dye Live Dead Near IR. Cells were washed in FACS buffer and assessed by flow cytometry. The frequency of cells falling with-in the viability gate is shown above the sample for each corresponding time-point.

### Flow cytometry panel design and representative experimental data

The antibody panel design is shown in **Table 1**. The panel was designed based on availability of pre-conjugated antibodies from commercial vendors against phosphorylated STAT proteins. The selected clones were tested for compatibility with the fixation/permeabilization protocol used and titrated to find the optimal staining concentration.

Plots of representative staining and the general gating strategy are shown in **Figure 2**. A forward-scatter vs side-scatter gate was initially used to gate on the cell populations, excluding debris (lower left corner) and DynaBeads (upper left corner). This was followed by forward-scatter height vs. area and side-scatter height vs. area gates to exclude doublets. Live Dead amine reactive dye negative cells (i.e. viable cells) were then gated.

**Figure 2.**
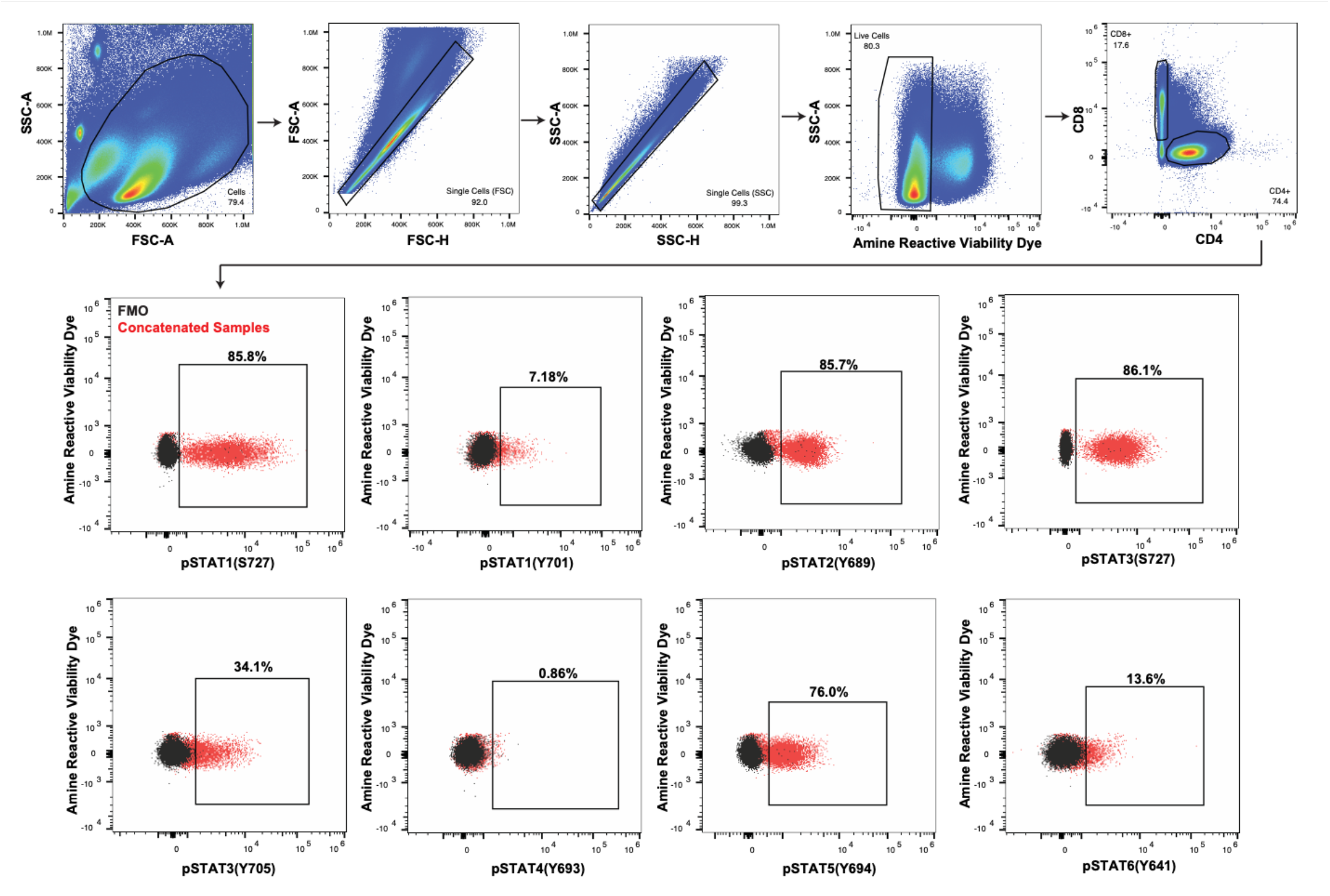
Flow cytometry gating strategy. T-cells were activated via CD3/CD28 stimulation for various durations and assessed by flow cytometry for expression of the indicated markers. The resulting fcs files concatenated. Debris were excluded using a FSC-A vs. SSC-A gate. Doublets were excluded using an FSC-H vs. FSC-A and subsequent SSC-H vs. SSC-A gate. Viable cells were gated based on absence of the amine reactive viability dye Live Dead Near IR. Viable cells were next gated into CD4+ and CD8+ populations. The plots in the second and third rows are from the CD4+ population. Expression of each pSTAT protein/residue in the concatenated samples is shown in red and the corresponding FMO in black. The indicated frequencies of positive cells in the concatenated samples are shown above the corresponding gate.

For most experiments, we enriched T-cells by a CD3 negative selection kit, so we did not include a CD3 antibody in our panel. In the plots show in in **Figure 2**, CD4+ cells were gated. Plots for each evaluated pSTAT are shown, with the concatenated samples in red and the corresponding fluorescence minus one (FMO) control sample in black.

### Activation by CD3/CD28 stimulation enhances expression of pSTATs

We initially tested our panel to assess the impact of polyclonal T-cell activation on the expression of pSTATs. Cells were left unstimulated (0h timepoint) or activated for times indicated in **Figure 3A**. We evaluated several methods of CD3 and CD28 polyclonal activation: 1) plate bound CD3 (clone OKT3, 0.5ug/mL) with soluble CD28 (clone CD28.1, 0.5ug/mL), 2) Immunocult (CD3 and CD28 antibody complexes), and 3) Dynabeads (CD3 and CD28 antibodies bound to magnetic beads) at a 1:1 (beads to cells) ratio or a 1:10 ratio. At baseline (i.e. no activation), high frequencies of pSTAT1(S727), pSTAT2(Y689) and pSTAT3(S727) expressing T-cells were observed. Both pSTAT1 (S727) and pSTAT3(S727) showed two positivity states, which we termed “low” and “high”. Gating of these markers into “low” and “high” populations showed that pSTAT1(S727)^low^ and pSTAT3(S727)^low^ frequencies were elevated at baseline, but the “high” states were absent. Increases in pSTAT1(S727)^high^, pSTAT1(Y701), pSTAT3(S727)^high^, pSTAT3(Y705), pSTAT4(Y693) and pSTAT6(Y641) occurred at the 24 hour timepoint. Only minor, inconsistent changes were observed in pSTAT5(Y694) expression. Representative histograms of pSTAT expression at the 72-hour time-point are shown in **Figure 3B**, with the inclusion of an FMO sample in solid grey.

**Figure 3.**
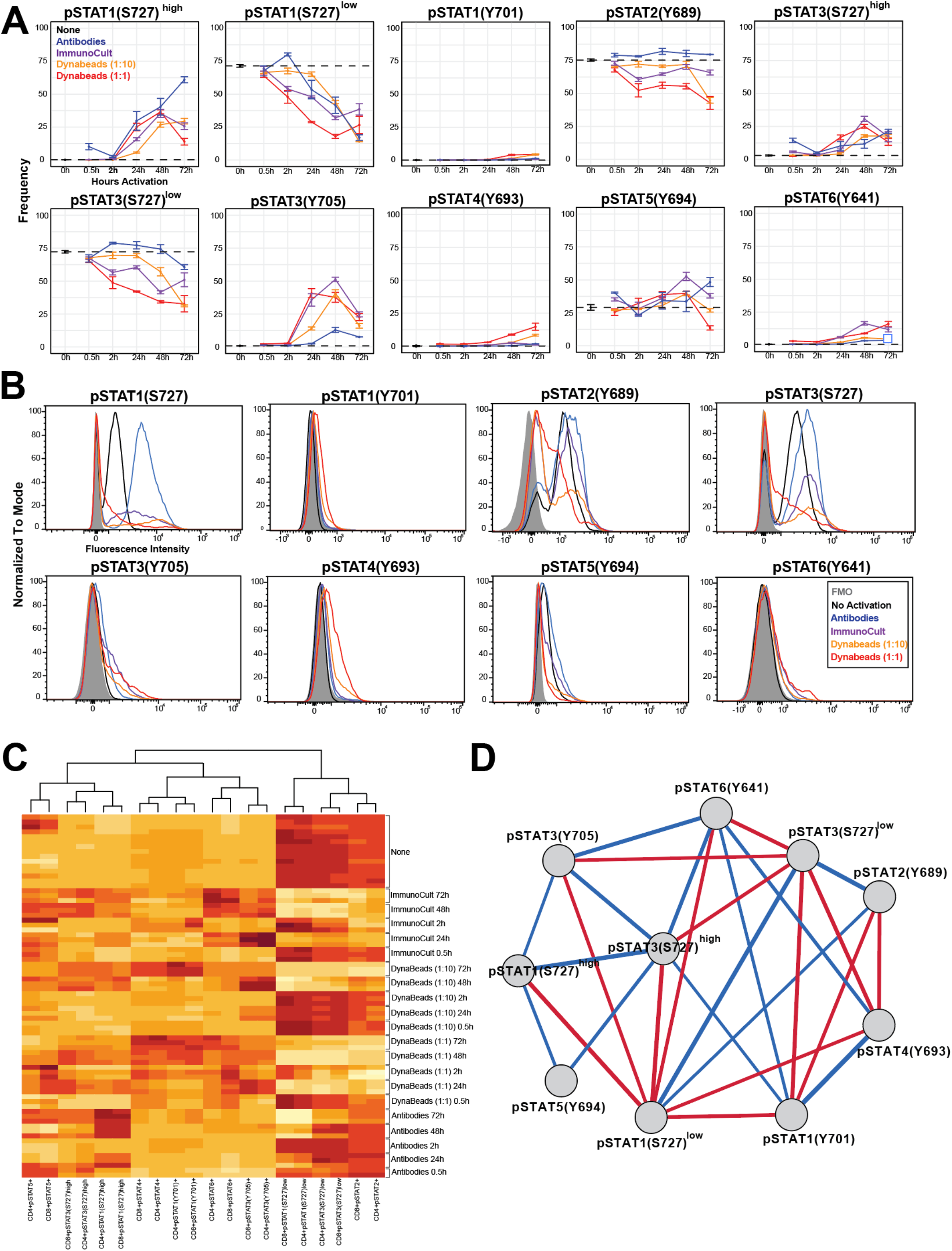
T-cell activation via CD3/CD28 stimulation enhances expression of all pSTATs. T-cells cultured with the indicated activating modalities for 15 minutes, 30 minutes, 24 hours, or 48 hours. Each condition and timing pairing was done in triplicate. (**A)** Each panel shows the kinetics of a different pSTAT residue. Cultures activated with CD3/CD28 DynaBeads at a 1:1 ratio are shown in red, DynaBeads at a 1:10 (bead to T-cell) ratio in orange, CD3/CD28 ImmunoCult activation in purple and plate bound CD3 antibodies with soluble CD28 antibodies in blue. Zero-hour values were obtained from non-activated T-cell cultures (black dotted lines). Error bars show the standard error of the mean (SEM). (**B)** Representative histograms for expression of the different pSTAT residues in CD4+ T-cells at the 72-hour timepoint are shown. Corresponding FMO samples are shown in shaded grey, non-activated T-cells in black, 1:10 ratio Dynabead activated T-cells in orange, and late bound CD3 antibodies with soluble CD28 antibodies in blue. (**C)** A heatmap of frequencies of pSTAT+ CD4+ and CD8+ T-cells is shown. Column data (pSTATs) were scaled and centered and ordered/grouped based on hierarchical clustering using Euclidean distance. (**D)** Frequencies of pSTAT+ CD4+ T-cells were assessed for correlations and visualized as a network. The Pearson correlation coefficient values are represented by the edges connecting parameter nodes, with positive correlations in blue and negative correlations in red. The thickness of the edges indicates the magnitude of the correlation. Edges were filtered based on the p-values associated to the correlations using p < 0.05 to provide a better visualization. Results shown are representative of two independent experiments.

We next evaluated the co-expression profiles of the evaluated pSTATs. A heatmap of pSTAT positivity frequencies across the different activation methods/samples is shown in **Figure 3C** and a network of correlations in **Figure 3D**. Pearson r-values are reported in **Supplemental Table 1** and the corresponding p-values **in Supplemental Table 2**. pSTAT expression profiles for CD4+ and CD8+ T-cells were highly similar and clustered together (**Figure 3C**). We observed several positive correlation clusters between pSTATs, including pSTAT2/pSTAT1(S727)^low^/pSTAT3(S727)^low^, pSTAT4/pSTAT1(Y701)/pSTAT6/pSTAT3(Y705), and pSTAT5/pSTAT3(S727)^high^/pSTAT1 (S727)^high^ (**Figure 3D**). The pSTAT2/pSTAT1 (S727)^low^/pSTAT3(S727)^low^ cluster was elevated in non-activated T-cells and negatively correlated with the other two clusters.

### Cytokine stimulation induces rapid and transient increases in pSTAT expression

We next assessed the impact of cytokine stimulation on T-cell pSTAT expression. In an initial experiment evaluating a panel of recombinant cytokines (data not shown), we found that IFNβ was a potent stimulator of all pSTATs. Therefore, we evaluated three concentrations of IFNβ (10, 50 and 200ng/mL) at different time points (0.25, 0.5, 1, 2 and 24 hours) for their impact on pSTAT induction in T-cells (**Figure 4A**). Increases in the frequency of T-cells expressing pSTAT1(Y701), pSTAT3(Y705) and pSTAT4(Y693) were observed as early as the 15-minute timepoint, but there were negligible differences observed at different concentrations of IFNβ. No changes were observed in frequencies of pSTAT2(Y689) expressing T-cells, which were ~90% positive at baseline. While inconsistent across timepoints, IFNβ decreased pSTAT1(S727)^high^ and pSTAT3(S727)^high^ frequencies while increasing pSTAT1(S727)^low^ and pSTAT3(S727)^low^ frequencies in a dose-dependent manner. Interestingly, pSTAT6(Y641) frequencies were increased by IFNβ stimulation, but in an inverse dose-dependent manner. However, the absolute frequency of pSTAT6 positive cells was low (~1.5% control and ~3% at 30-minutes in the 10ng/mL group).

**Figure 4.**
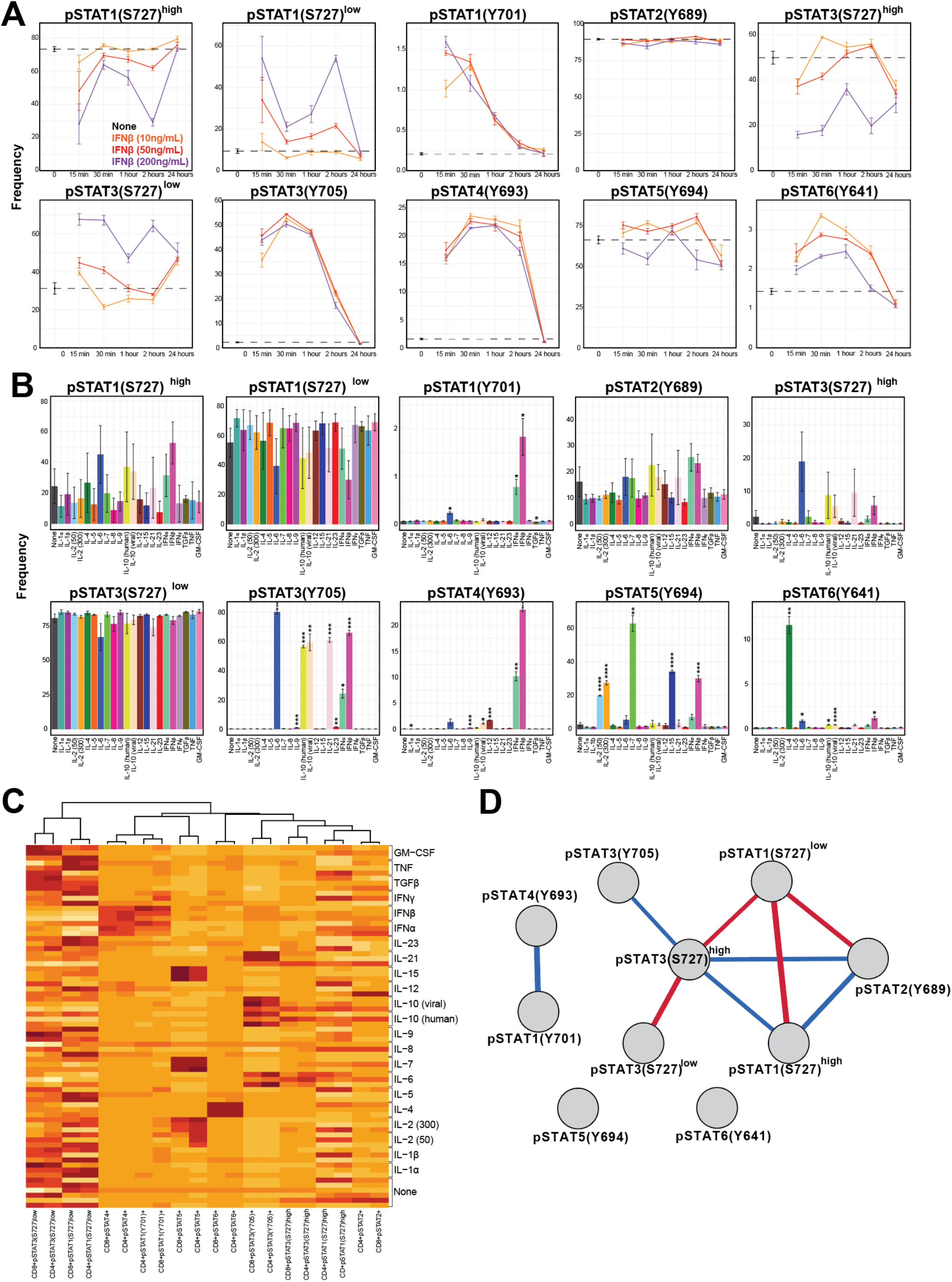
Assessment of pSTAT induction by a panel of recombinant cytokines. (**A)** T-cells were isolated via negative CD3+ selection and cultured with IFNβ in triplicate for the indicated amounts of time. The untreated (“none”) group frequencies are shown at the 0 time-point and by the dotted black line. The IFNβ concentrations are shown as follows: 10ng/mL in orange, 50ng/mL in red, and 200ng/mL in purple. Error bars show the SEM. (**B)** T-cells were isolated via negative CD3+ selection and cultured with the indicated recombinant cytokines for 30 minutes prior to assessment using the flow cytometry protocol. Each was done in triplicate, apart from the untreated (“none”) group which had six replicates. Each panel shows the frequency of positive CD4+ T-cells for the different pSTAT residues. Error bars show the SEM. Significance was determined by a Bonferroni corrected t-tests. Statistical significance is indicated as follows: * p<0.05, ** p<0.01, ***p<0.001, ****p<0.0001. (**C)** A heatmap of frequencies of pSTAT+ CD4+ and CD8+ T-cells is shown. Column data (pSTATs) were scaled and centered and ordered/grouped based on hierarchical clustering. (**D)** Frequencies of pSTAT+ CD4+ T-cells were assessed for correlations and visualized as a network. The Pearson correlation coefficient values are represented by the edges connecting parameter nodes, with positive correlations in blue and negative correlations in red. The thickness of the edges indicates the magnitude of the correlation. Edges were filtered based on the p-values associated to the correlations using p < 0.05 to provide a better visualization. Results shown are representative of two independent experiments.

### Evaluation of a panel of recombinant cytokines on induction of pSTATs

We choose the 30-minute timepoint and 50ng/mL (or international units (IU) in the case of IL-2) concentration to next evaluate a panel of 21 recombinant cytokines. We also included an additional 300IU IL-2 treatment group given that different concentrations of IL-2 differentially impact effector vs. Tregs^41^. As before, IFNβ increased the frequency of expressing T-cell for all pSTATs, with the exception of pSTAT1(S727)^low^ and pSTAT3(S727)^low^ (**Figure 4B**). IFNα also increased the frequency T-cells expressing these pSTATs, except for pSTAT6(Y641). Canonical associations were also observed including: IL-6/IL-10/IL-21 induction of pSTAT3(Y705), IL-2/IL-7/IL-15 induction of pSTAT5(Y694) and IL-4 induction of pSTAT6(Y641). Interestingly, IL-6 significantly increased the frequencies of pSTAT1(Y701), pSTAT3(Y705), pSTAT6(Y641) and trended towards increases in pSTAT3(S727)^high^ and pSTAT4(Y693).

We also evaluated the co-expression profiles of pSTATs by T-cells stimulated with cytokines. A heatmap of pSTAT positivity frequencies across the different activation methods/samples is shown in **Figure 4C**, and a network of correlations in **Figure 4D**. Pearson r-values are reported in **Supplemental Table 3** and the corresponding p-values **in Supplemental Table 4**. As with activated T-cells, CD4+ and CD8+ T-cells had similar pSTAT expression profiles in response to different cytokine stimulations and clustered together. Likewise, pSTAT2/pSTAT1(S727)^low^/pSTAT3(S727)^low^ expression was positively correlated, as was pSTAT4/pSTAT1(Y701).

### *In vitro* polarization of T-cells results in complex pSTAT signaling

Finally, we assessed changes in pSTAT signaling resulting from polarization. Naïve CD4+ T-cells were cultured in polarizing conditions for 30 minutes, 24 hours or 96 hours and then evaluated for expression of pSTATs. A portion of cells were used to validate polarization by a separate intracellular flow cytometry panel (**Supplemental Figure 1A**). Th1 polarized T-cells had the highest frequency of IFNγ expression, Th2 polarized cells had the highest frequency of IL-4, and iTreg polarized cells had the highest frequency of FOXP3 expression. Frequencies of canonically associated pSTATs were upregulated in polarized cells: pSTAT1(Y701) in Th1, pSTAT6(Y641) in Th2, pSTAT3(Y705) in Th17, and pSTAT5(Y694) in iTregs (**Figure 5A and 5B**). Th1 and iTregs also had increases in pSTAT1(S727)^high^ expression. Th1 polarized also had increases in pSTAT3(Y705) and pSTAT4(Y649) frequencies.

**Figure 5.**
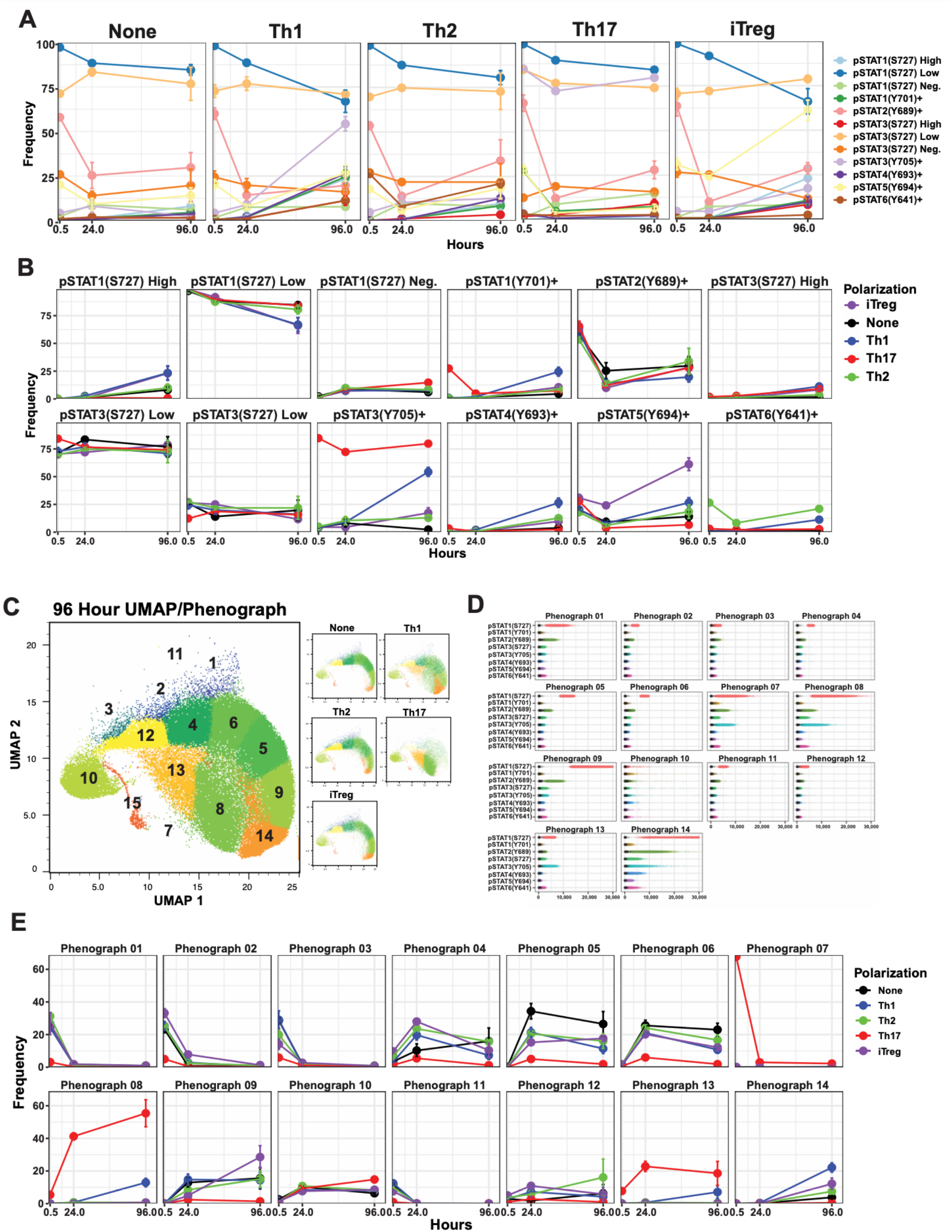
*In vitro* polarization of naïve CD4+ T-cells results in complex pSTAT expression profiles. Naïve CD4+ T-cells were isolated and polarized as indicated in the methods section. Polarizations were done in triplicates. Cells were assessed for the expression of pSTATs after 30 minutes, 24 hours and 96 hours of polarization. (**A)** Each panel shows the expression of the pSTATs in the different polarization groups at the different time points. (**B)** Each panel shows the polarization frequencies for each pSTAT at different time points. (**C)** The eight pSTAT parameters were reduced to two dimensions by UMAP and assessed for clusters by Phenograph. The concatenated UMAP of the 96-hour timepoints is shown in the large panel, with the Phenograph clusters labeled by number. The 96-hour UMAPs of each polarization group are shown in the smaller panels. (**D)** The expression profiles for each pSTAT in each of the Phenograph clusters (separate panels) is plotted. Each cell belonging to the corresponding Phenograph cluster is plotted with its value on the x-axis representing the MFI value for the corresponding y-axis parameter. The MFI values for the corresponding FMO for each pSTAT are shown as black dots. (**E)** The frequencies of the polarization groups in each Phenograph cluster are plotted. Results shown are representative of two independent experiments.

To further investigate the complex pSTAT expression profiles in polarized T-cells, we performed dimension reduction by UMAP and clustering by Phenograph. The UMAP colored by phonograph clusters for all samples and by time-point is shown in **Supplemental Figure 1B**. At 96 hours, clustering for Th1 and Th17 cells were distinct, but Th2 and iTregs were similar to the unpolarized “None” group (**Figure 5C**). Th1 polarized cells had high frequencies of cluster 14, Th2 polarized cells had high frequencies of cluster 12, Th17 polarized cells had high frequencies of clusters 8 and 13, and iTregs had high frequencies of cluster 9 (**Figure 5E**). To evaluate the phenotypes associated with each cluster, we plotted the expression of pSTATs for each cell in the cluster, overlaying the corresponding FMO events to mask background from positive signal (**Figure 5D**). Cluster 14 (Th1) had high expression of all the pSTATs evaluated. Cluster 12 (Th2) was mostly absent of expression of all pSTATs. Cluster 8 (Th17) had relatively high expression of pSTAT3(Y705) and cluster 13 (Th17) had low expression of pSTAT1(S727) and moderate expression of pSTAT3(Y705). Cluster 9 (iTreg) had high expression of pSTAT1(S727) and relatively high expression of pSTAT5(Y694).

## Discussion

We have developed and optimized a sensitive methodology for simultaneously assessing tyrosine phosphorylation on six STAT proteins and additional serine phosphorylation on pSTAT1 and pSTAT3, at single cell resolution. This approach is optimized for a 96-well format, making it relatively rapid and compatible with medium throughput assays. While we have utilized this approach to interrogate T-cell biology in this study, it is readily compatible with all human cell types. These characteristics make this a potentially powerful tool in interrogating the roles of STAT signaling in health and disease.

To demonstrate the utility of this panel, we interrogated the impact of activation, cytokine stimulation and polarization on T-cell pSTAT signaling. In doing so, we observed several novel findings. This includes the observation that pSTAT2(Y689) is constitutively expressed in most T-cells, irrespective of activation or stimulation. Previous studies have shown that STAT2 phosphorylation is driven by type I IFNs^42^, but how and if may be related to the observed constitutive expression of pSTAT2 in T-cells is unclear. We also observed that IFNβ and, to a lesser degree, IFNα, rapidly increased the frequency of expression of all pSTATs. Again, the biology underlying this remains unclear.

We observed that like pSTAT2, both pSTAT1(S727) and pSTAT3(S727) were expressed in most T-cells. We defined two positivity states for each of these markers, “low” and “high”. Activation and specific cytokine stimulation caused shifts between the low to high population. In contrast, pSTAT1(Y701) and pSTAT3(Y705) were absent in resting cells. This contrast suggests that phosphorylation of serine and the tyrosine residues have distinct roles in T-cell biology. It also raises the question of whether similar differences exist in other STAT proteins. Due to the lack of commercially available antibodies, we did not investigate phosphorylation of serine residue in other STAT molecules. We anticipate that as higher dimension flow cytometry becomes increasingly common and additional antibodies are made available, future refinement of this panel will be able to incorporate additional phosphorylated residues.

The results of this study highlight that the often-used model of one pSTAT induced by cytokine stimulation (e.g. IL-10 induction of pSTAT3) or associated with T-cell polarization (e.g. pSTAT5 in Tregs) is oversimplified. For example, the data in this study demonstrated that IL-10 induces not only pSTAT3 signaling, but also pSTAT6 and possibly pSTAT2. In the case of polarization, the data in this study demonstrated that pSTAT5 was upregulated in iTregs as well as Th1 cells, and that Th1 polarization was accompanied by high frequencies of pSTAT3(Y705) expressing T-cells. We hypothesize that formation of homo- and heterodimers and competition between these different pSTAT complexes add another layer of complexity that also needs to be assessed in studies of pSTAT signaling. Computational modeling supports this hypothesis^43^. Though we did not test it in the present study, this panel is theoretically compatible with imaging cytometry, which would allow for assessing dimerization by co-localization, as well as cellular location (i.e. cytoplasmic or nuclear).

Collectively, the presented study details a flow cytometry-based protocol for the simultaneous assessment of eight phosphorylated residues on six STAT proteins. This protocol has the potential to be a useful tool in future investigations of the roles of STAT signaling in T-cells and other cell types in health and disease.

## Author Contributions

- Emily Monk: analyzed experiments, assisted in manuscript assembly, wrote data analysis code.
- Melinda Vassallo: conducted experiments.
- Paulo Burke: analyzed experiments, assisted in manuscript assembly, wrote data analysis code.
- Jeffrey Weber: assisted in manuscript assembly and editing.
- Pratip Chattopadhyay: reviewed data, assisted with manuscript assembly.
- David Woods: designed and optimized panel; designed, conducted, and analyzed experiments; assisted in manuscript assembly; wrote data analysis code.

## Acknowledgements

We extend our appreciation to Ann Strange for her assistance in creating the Docker container and to Carol Amato and Brian Thompson for their assistance in manuscript editing.

## Supplemental Tables

**Supplemental Table 1.**
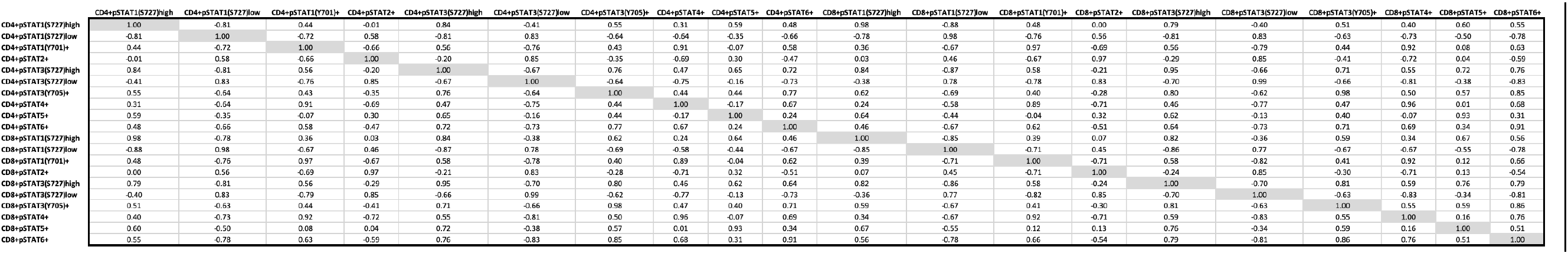
Pearson R-values for pSTATs in activated T-cells.

**Supplemental Table 2.**
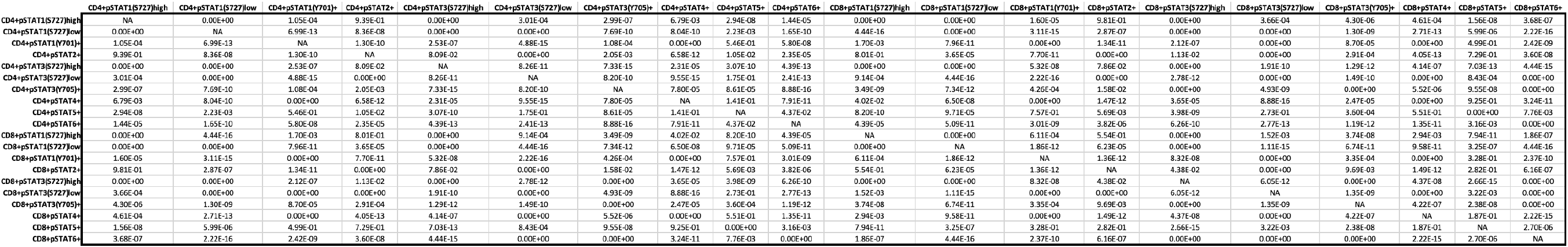
P-values for activated T-cell pSTAT correlations.

**Supplemental Table 3.**
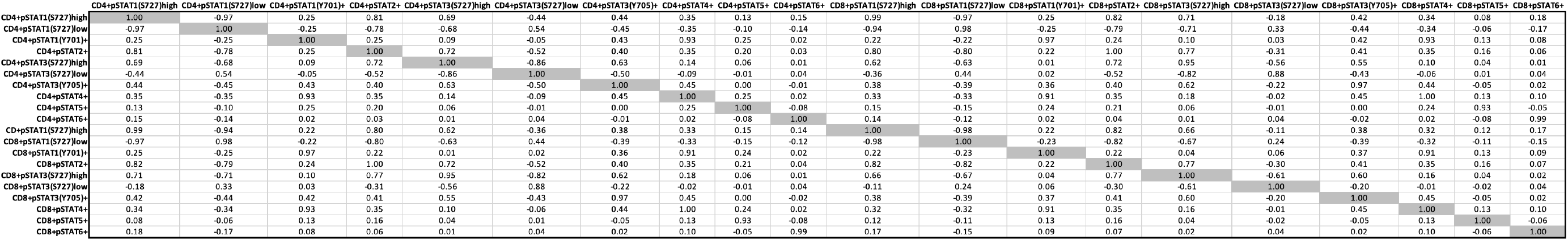
Pearson R-values for pSTATs in cytokine stimulated T-cells.

**Supplemental Table 4.**
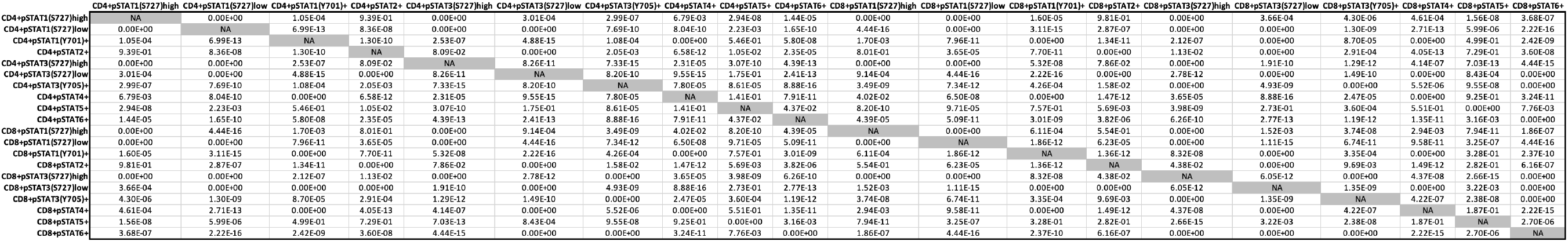
P-values for cytokine stimulated T-cell pSTAT correlations.

**Supplemental Figure 1.**
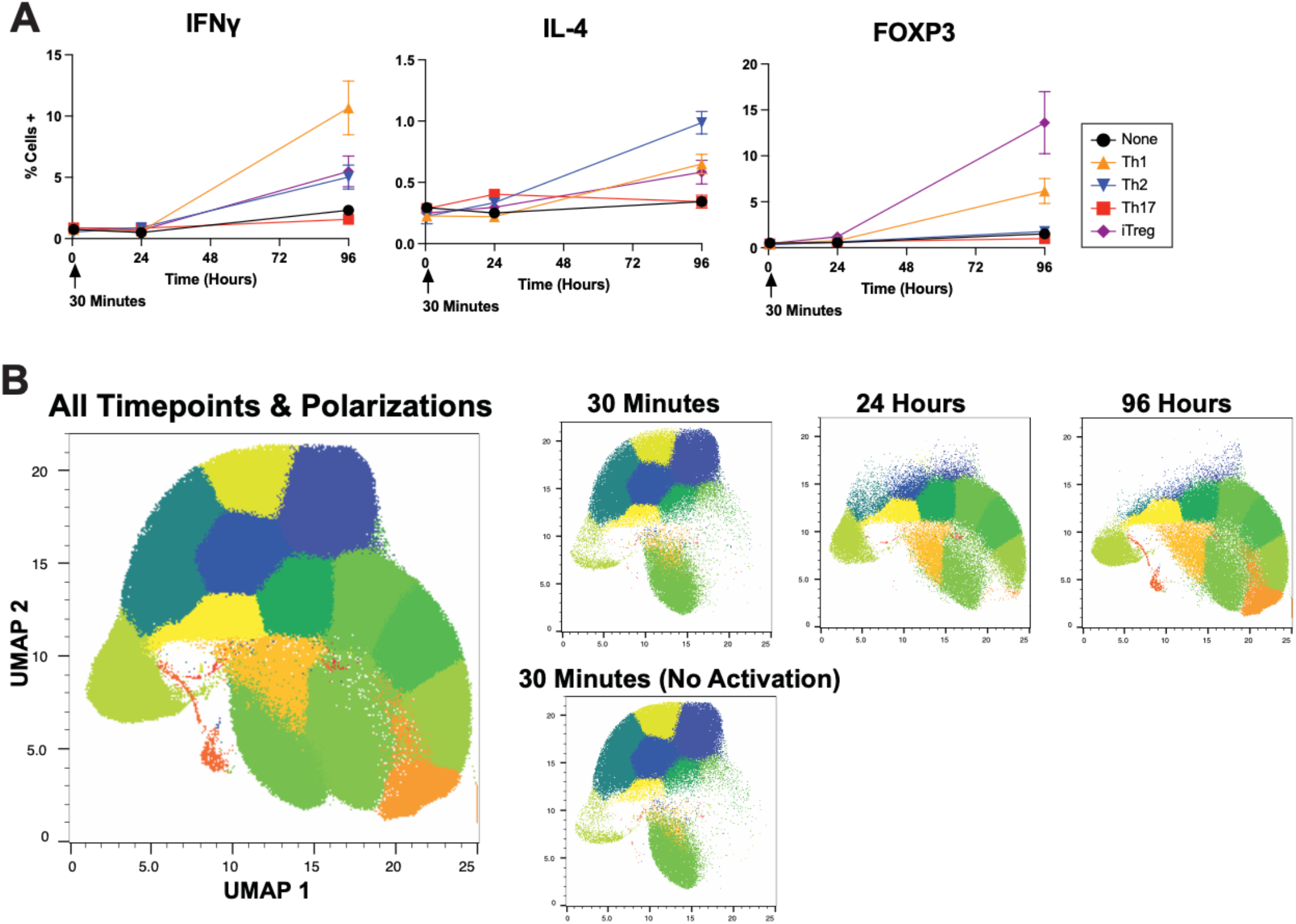
Polarization of T-cells. Naïve CD4+ T-cells were cultured in different polarizing conditions. (**A)** Cultures were evaluated by intracellular flow cytometry at the indicated time points for frequencies of IFNγ (leftmost panel), IL-4 (middle panel) and FOXP3 (rightmost panel) expressing cells. The “None” group (i.e. cells with only CD3/CD28 activation) are shown in black, the Th1 polarization samples in orange, Th2 polarized samples in blue, Th17 polarized samples in red, and iTreg polarized samples in purple. Polarizations were performed in triplicate. (**B)** All timepoints and polarizations were concatenated and reduced to two dimensions by UMAP. Phenograph clustering was overlayed as a third, color dimension. Subplots of each timepoint are also shown. Results shown are representative of two independent experiments.

## References

1 Aaronson, D. S. & Horvath, C. M. A road map for those who don’t know JAK-STAT. Science 296, 1653–1655, doi:10.1126/science.1071545 (2002).

2 Grimley, P. M., Dong, F. & Rui, H. Stat5a and Stat5b: fraternal twins of signal transduction and transcriptional activation. Cytokine Growth Factor Rev 10, 131–157, doi:10.1016/s1359-6101(99)00011-8 (1999).

3 Lin, J. X. & Leonard, W. J. The role of Stat5a and Stat5b in signaling by IL-2 family cytokines. Oncogene 19, 2566–2576, doi:10.1038/sj.onc.1203523 (2000).

4 Jenks, J. A. et al. Differentiating the roles of STAT5B and STAT5A in human CD4+ T cells. Clin Immunol 148, 227–236, doi:10.1016/j.clim.2013.04.014 (2013).

5 Seif, F. et al. The role of JAK-STAT signaling pathway and its regulators in the fate of T helper cells. Cell Commun Signal 15, 23, doi:10.1186/s12964-017-0177-y (2017).

6 Siegel, A. M. et al. A critical role for STAT3 transcription factor signaling in the development and maintenance of human T cell memory. Immunity 35, 806–818, doi:10.1016/j.immuni.2011.09.016 (2011).

7 Kanai, T. et al. Identification of STAT5A and STAT5B target genes in human T cells. PLoS One 9, e86790, doi:10.1371/journal.pone.0086790 (2014).

8 Harrington, L. E. et al. Interleukin 17-producing CD4+ effector T cells develop via a lineage distinct from the T helper type 1 and 2 lineages. Nat Immunol 6, 1123–1132, doi:10.1038/ni1254 (2005).

9 Durant, L. et al. Diverse targets of the transcription factor STAT3 contribute to T cell pathogenicity and homeostasis. Immunity 32, 605–615, doi:10.1016/j.immuni.2010.05.003 (2010).

10 Betts, B. C., Veerapathran, A., Pidala, J., Yu, X. Z. & Anasetti, C. STAT5 polarization promotes iTregs and suppresses human T-cell alloresponses while preserving CTL capacity. J Leukoc Biol 95, 205–213, doi:10.1189/jlb.0313154 (2014).

11 Kaplan, M. H., Schindler, U., Smiley, S. T. & Grusby, M. J. Stat6 is required for mediating responses to IL-4 and for development of Th2 cells. Immunity 4, 313–319, doi:10.1016/s1074-7613(00)80439-2 (1996).

12 Owen, K. L., Brockwell, N. K. & Parker, B. S. JAK-STAT Signaling: A Double-Edged Sword of Immune Regulation and Cancer Progression. Cancers (Basel) 11, doi:10.3390/cancers11122002 (2019).

13 Woods, D. M. et al. Decreased Suppression and Increased Phosphorylated STAT3 in Regulatory T Cells are Associated with Benefit from Adjuvant PD-1 Blockade in Resected Metastatic Melanoma. Clin Cancer Res 24, 6236–6247, doi:10.1158/1078-0432.CCR-18-1100 (2018).

14 Wang, Y., Shen, Y., Wang, S., Shen, Q. & Zhou, X. The role of STAT3 in leading the crosstalk between human cancers and the immune system. Cancer Lett 415, 117–128, doi:10.1016/j.canlet.2017.12.003 (2018).

15 Verhoeven, Y. et al. The potential and controversy of targeting STAT family members in cancer. Semin Cancer Biol 60, 41–56, doi:10.1016/j.semcancer.2019.10.002 (2020).

16 Schultz, J. et al. Tumor-promoting role of signal transducer and activator of transcription (Stat)1 in late-stage melanoma growth. Clin Exp Metastasis 27, 133–140, doi:10.1007/s10585-010-9310-7 (2010).

17 Song, L., Turkson, J., Karras, J. G., Jove, R. & Haura, E. B. Activation of Stat3 by receptor tyrosine kinases and cytokines regulates survival in human non-small cell carcinoma cells. Oncogene 22, 4150–4165, doi:10.1038/sj.onc.1206479 (2003).

18 Xu, Y. H. & Lu, S. A meta-analysis of STAT3 and phospho-STAT3 expression and survival of patients with non-small-cell lung cancer. Eur J Surg Oncol 40, 311–317, doi:10.1016/j.ejso.2013.11.012 (2014).

19 Chiba, T. STAT3 Inhibitors for Cancer Therapy –the Rationale and Remained Problems. EC Cancer 1, S1–S8 (2016).

20 Kortylewski, M., Jove, R. & Yu, H. Targeting STAT3 affects melanoma on multiple fronts. Cancer Metastasis Rev 24, 315–327, doi:10.1007/s10555-005-1580-1 (2005).

21 Kalbasi, A. & Ribas, A. Tumour-intrinsic resistance to immune checkpoint blockade. Nat Rev Immunol 20, 25–39, doi:10.1038/s41577-019-0218-4 (2020).

22 Pfitzner, E., Kliem, S., Baus, D. & Litterst, C. M. The role of STATs in inflammation and inflammatory diseases. Curr Pharm Des 10, 2839–2850, doi:10.2174/1381612043383638 (2004).

23 O’Shea, J. J. & Plenge, R. JAK and STAT signaling molecules in immunoregulation and immune-mediated disease. Immunity 36, 542–550, doi:10.1016/j.immuni.2012.03.014 (2012).

24 Warshauer, J. T. et al. A human mutation in STAT3 promotes type 1 diabetes through a defect in CD8+ T cell tolerance. J Exp Med 218, doi:10.1084/jem.20210759 (2021).

25 Jakkula, E. et al. Genome-wide association study in a high-risk isolate for multiple sclerosis reveals associated variants in STAT3 gene. Am J Hum Genet 86, 285–291, doi:10.1016/j.ajhg.2010.01.017 (2010).

26 Remmers, E. F. et al. STAT4 and the risk of rheumatoid arthritis and systemic lupus erythematosus. N Engl J Med 357, 977–986, doi:10.1056/NEJMoa073003 (2007).

27 Wen, Z., Zhong, Z. & Darnell, J. E., Jr. Maximal activation of transcription by Stat1 and Stat3 requires both tyrosine and serine phosphorylation. Cell 82, 241–250, doi:10.1016/0092-8674(95)90311-9 (1995).

28 Zhu, X., Wen, Z., Xu, L. Z. & Darnell, J. E., Jr. Stat1 serine phosphorylation occurs independently of tyrosine phosphorylation and requires an activated Jak2 kinase. Mol Cell Biol 17, 6618–6623, doi:10.1128/MCB.17.11.6618 (1997).

29 Huang, G., Yan, H., Ye, S., Tong, C. & Ying, Q. L. STAT3 phosphorylation at tyrosine 705 and serine 727 differentially regulates mouse ESC fates. Stem Cells 32, 1149–1160, doi:10.1002/stem.1609 (2014).

30 Levine, J. H. et al. Data-Driven Phenotypic Dissection of AML Reveals Progenitor-like Cells that Correlate with Prognosis. Cell 162, 184–197, doi:10.1016/j.cell.2015.05.047 (2015).

31 Leland McInnes, J. H., James Melville. UMAP: Uniform Manifold Approximation and Projection for Dimension Reduction. arXiv (2020).

32 R Core Team. R: A language and environment for statistical computing, <https://www.R-project.org/> (2020).

33 Ellis B, H. P., Hahne F, Le Meur N, Gopalakrishnan N, Spidlen J, Jiang M, Finak G. flowCore: flowCore: Basic structures for flow cytometry data. R package version 2.6.0 (2021).

34 Hadley Wickham, M. A., Jennifer Bryan, Winston Chang, Lucy D’Agostino McGowan, Romain Francois, Garrett Grolemund, Alex Hayes, Lionel Henry, Jim Hester, Max Kuhn, Thomas Lin Pedersen, Evan Miller, Stephan Milton Bache, Kirill Muller, Jeroen Ooms, David Robinson, Dana Paige Seidel, Vitalie Spinu, Kohske Takahashi, David Vaughan, Claus Wilke, Kara Woo, Hiroaki Yutani. Welcome to the Tidyverse. The Journal of Open Source Software 4, 1686, doi:https://doi.org/10.21105/joss.01686 (2019).

35 Wickham, H. ggplot2: Elegant Graphics for Data Analysis. Springer-Verlag (2016).

36 Coombes KR, B. G., Abrams ZB, Abruzzo LV Polychrome: Creating and Assessing Qualitative Palettes with Many Colors. Journal of Statistical Software 90, 1–23, doi:10.18637/jss.v090.c01 (2019).

37 Kassambara, A. ggpubr: ‘ggplot2’ Based Publication Ready Plots. (2020).

38 Jr., F. E. H. Hmisc: Harrell Miscellaneous. (2021).

39 Csardi G, N. T. The igraph software package for complex network research. InterJournal, Complex Systems 1695(2006).

40 Shannon, P. et al. Cytoscape: a software environment for integrated models of biomolecular interaction networks. Genome Res 13, 2498–2504, doi:10.1101/gr.1239303 (2003).

41 Liao, W., Lin, J. X. & Leonard, W. J. Interleukin-2 at the crossroads of effector responses, tolerance, and immunotherapy. Immunity 38, 13–25, doi:10.1016/j.immuni.2013.01.004 (2013).

42 Park, C., Li, S., Cha, E. & Schindler, C. Immune response in Stat2 knockout mice. Immunity 13, 795–804, doi:10.1016/s1074-7613(00)00077-7 (2000).

43 Sadreev, II et al. The competitive nature of STAT complex formation drives phenotype switching of T cells. Immunology, doi:10.1111/imm.12851 (2017).

