## Supplemental Methods for "Simultaneous assessment of eight phosphorylated STAT residues in T-cells by flow cytometry"

Phosphorylated STAT flow cytometry staining protocol (96 well format).

1. Move Perm Buffer III to -20C freezer.
2. Perform treatments and timepoint according to experiment.
3. For the last five minutes of culture add 50uL of 5X (5uL per 1mL) LD Near IR to all wells (based on 200uL culture volume).
4. Centrifuge at 4C for 3 minutes. All centrifugation for this protocol needs to be at 4 degrees Celsius.
5. Remove media by normal “dumping” in sink.
6. Add 300uL per well of cold 1X TFP Fix/Perm buffer. *Be sure to make all buffers prior and to keep them cold.
7. Put plate in 4C fridge for 50 minutes.
8. Centrifuge at 4C for 5 minutes. Dump.
9. Add 200uL per well of cold 1X TFP Perm/Wash buffer.
10. Repeat centrifugation. Dump.
11. Use vortex to break cell pellets.
12. Add 300uL per well of Perm buffer III.
13. Leave on ice for 20 minutes.
14. Centrifuge. Dump.
15. Add 200uL of 1X TFP Perm/Wash buffer.
16. Centrifuge. Dump.
17. Add 200uL of 1X TFP Perm/Wash buffer.
18. Centrifuge. Dump.
19. Add 30uL per well of 1X TFP containing antibody cocktail for staining.
20. Leave in 4C fridge for 40 minutes.
21. Centrifuge. Dump.
22. Add 200uL of 1X TFP Perm/Wash buffer.
23. Centrifuge. Dump.
24. Add 200uL of 1X TFP Perm/Wash buffer.
25. Centrifuge. Dump.
26. Resuspended in 200uL per well of cold FACS buffer.
27. Run flow. *if not able to run immediately, keep in 4C until running.
